# Opposing mechanical anchorage drives collective cell–matrix interactions

**DOI:** 10.64898/2026.07.24.740441

**Authors:** Umnia Doha, Ahmadreza Kashefi, William Drennan, M Taher A Saif

## Abstract

Collective cell behaviors emerge from mechanical interactions with the extracellular matrix (ECM), yet the physical principles governing long-range cell–cell communication remain elusive. Existing models assume that neighboring cells couple by strain-stiffening the ECM between them, amplifying contractility through positive feedback. Here we show that pairwise interactions are insufficient. Instead, stable mechanical communication requires opposing mechanical anchors that allow a cell to strain-stiffen the matrix on both sides. Combining ECM strain mapping, direct cell-force measurements, and live-cell imaging, we find that isolated cell pairs generate only weak, stochastic matrix strains without persistent interactions. In contrast, cells supported by opposing neighbors, or rigid beads acting as mechanical anchors, generate large bilateral matrix strains, increase effective matrix stiffness, align collagen fibers, and form stable multicellular networks. To explain these observations, we develop a predictive mechanosensitive theory introducing effective matrix stiffness and a critical contractile force governing the transition from stochastic to persistent interaction. The theory predicts, and experiments confirm, that opposing mechanical anchorage enables cells to exceed the critical force, trigger collective matrix remodeling, and compact the matrix through collagen-fiber buckling. Together, these findings provide a unifying framework for understanding collective force generation in development, wound repair, fibrosis, and tumor progression.

## Main text

Collective social behaviors often emerge from communications and interactions between many individual players, such as birds flock(1), fish school(2), and ants colonize(3). In a similar vein, cell-cell communication and cell-extracellular matrix (ECM) interaction within living organisms direct collective cell behaviors, generating distinctive phenotypes on both single-cell (cell shape and polarization(4, 5)) and multicellular (long-range pattern and organization(6)) levels with diverse functions. Though these morphogenetic processes critically underlie development(7), wound healing(8, 9) and repair by epithelial cells acting in concert(10), and disease progression such as tumorigenesis(11, 12), the biophysical mechanisms that underlie such processes remain unclear: how multicellular collective behavior emerges, is there a unifying principle, what is the role of ECM mechanics in cell-cell communication over long distances, and by what mechanism?

A simple model for studying the emergence of collective cell behavior is the compaction of extracellular matrix (ECM) gels by living cells(13–15). Cells compact fibrous ECM only above a critical density, implying that collective mechanical interactions emerge over a characteristic cell–cell distance (16). For NIH 3T3 fibroblasts in collagen-I, this distance is ∼150 µm but increases to ∼250 µm when cell-sized rigid beads are introduced into the matrix, suggesting that the interactions leading to compaction are primarily mechanical (17).

Current models describe these interactions as pairwise: two neighboring cells strain-stiffen the ECM between them, increasing matrix stiffness, promoting cell polarization and contractility, and reinforcing this response through a positive mechanical feedback loop (18–23). However, experiments reveal a paradox. In low-cell-density collagen, neighboring cells frequently approach one another yet separate without forming stable interactions or compacting the matrix. Thus, proximity alone is insufficient. What, then, are the necessary and sufficient conditions for collective mechanical interaction?

Here, we propose a mechanosensitive theory that resolves this paradox by introducing (1) effective matrix stiffness and (2) a critical contractile force that is intrinsic to a cell type. The critical force governs the transition from stochastic to persistent state of the cell. A cell enters a positive feedback loop only if it can strain-stiffen the ECM on **both** sides, thereby increasing the effective stiffness (the mechanical resistance experienced by the contracting cell) that it senses, and its contractility exceeds the critical force. This bilateral mechanical anchorage requires a minimum of three mechanically coupled cells, providing a unifying physical principle for collective cell–ECM interactions and matrix compaction.

### Hypothesis on cell-ECM-cell interaction and gel compaction

Fibrous extracellular matrices (ECMs) exhibit nonlinear strain stiffening: tensile strain aligns collagen fibers, increasing stiffness along the direction of loading (24–26). Mechanosensitive cells exploit this property by pulling on ECM fibers and responding to the resulting effective matrix stiffness (27–29). In contrast, linear elastic matrices do not strain-stiffen and therefore cannot sustain this mechanical feedback.

We hypothesize that a cell enters a positive force–strain–force feedback loop in a soft matrix (19) only if it can strain-stiffen the ECM on **both** sides simultaneously, i.e., increase the effective stiffness of the matrix locally, and if its contractile force exceeds an intrinsic critical force. Neighboring cells stiffer than the matrix (30), or other rigid objects attached to the ECM, can serve as opposing mechanical anchors that support bilateral strain stiffening. An anchor on only one side is insufficient because the effective stiffness sensed by the cell remains limited by the softer side of the matrix (SI Fig. S1 with a simple example). Consequently, two isolated neighboring cells cannot sustain mechanical coupling, whereas a minimum of three mechanically coupled cells can form a closed tensile network in which each cell is supported by opposing anchors. A linear elastic matrix does not strain stiffen and hence may not support the tensile network.

Formation of this tensile network is necessary but not sufficient for macroscopic gel compaction. As contractile forces increase, the network compresses the enclosed ECM. Once the compressive stress exceeds the buckling threshold of collagen fibers, the matrix collapses, expelling interstitial fluid and producing rapid gel compaction.

We first test this hypothesis experimentally and then develop a predictive mechanosensitive theory that explains the observations and generalizes them across matrix stiffnesses, cell densities, and mechanical boundary conditions.

## Results

### Cells do not exhibit any collective mechanical interactions in a linear elastic matrix (colTgel)

To determine whether nonlinear matrix mechanics are required for collective behavior, we formed free-floating discs of ColT gel embedded with NIH 3T3 fibroblasts (2 M/mL). ColT gel is a linear elastic matrix with tunable stiffness (0.5 or 10 kPa; Methods). Regardless of stiffness, the gels showed no detectable compaction over 9 days (Fig. S2A,B). In contrast, fibroblasts at same density (2 M/ml) compacted fibrous collagen-1 (2 mg/mL, 0.5 kPa) discs to ∼20% of their initial volume within 12 h(17).

Time-lapse imaging revealed that cells in ColT gels extended filopodia, polarized, and migrated randomly (Fig. S2C,D; SI Movie 1) without forming the interconnected networks observed in collagen. Filopodia from neighboring cells occasionally approached one another but failed to establish persistent interactions. Tracking embedded 2-µm tracer beads showed that the strain between neighboring cells fluctuated throughout the experiment (Fig. S2E), reflecting stochastic filopodial extension and retraction without mechanical coupling through the matrix. Unlike fibrous collagen(31, 32), the linear elastic matrix therefore failed to support long-range mechanical communication. Consequently, cells did not enter the positive mechanical feedback loop or form the tensile network required for matrix compaction.

### Pairwise interactions are insufficient for matrix stiffening in fibrous collagen

At low cell density, fibroblasts embedded in collagen-1 (2 mg/ml) do not compact the gel but migrate randomly(17). Neighboring cells occasionally approach one another stochastically, but subsequently separate without establishing stable interactions (Fig. 1A,B; SI Movie 2), resembling their behavior in linear elastic ColT gels.

**Fig 1.**
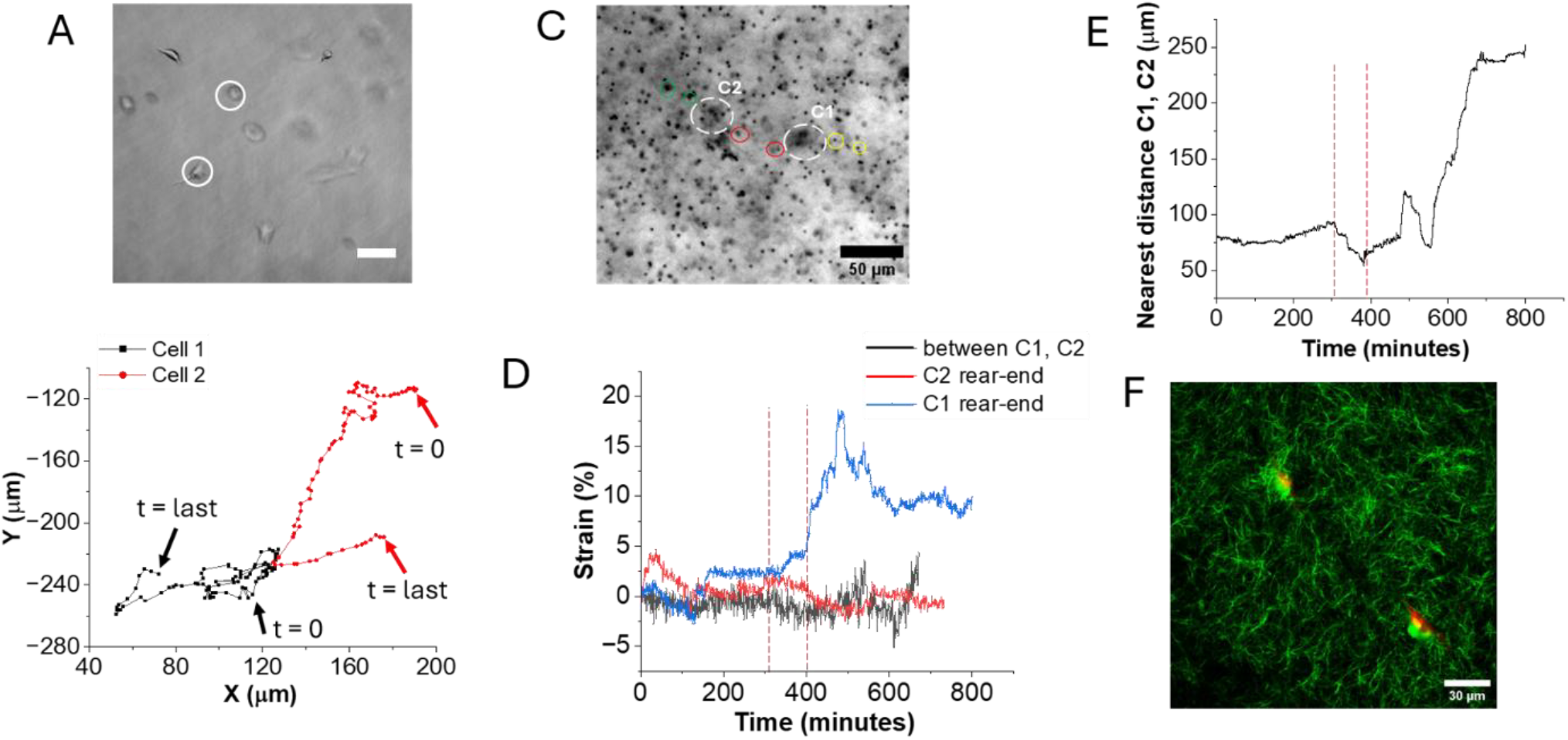
Fibroblasts (3T3) in collegen-1 matrix at low density (0.1 M cells/ml) do not interact with one another. (A) Brightfield image of 2 cells in a collagen disc tracked for time-lapse imaging. (B) Trajectory of the cells in (A) showing Brownian dynamics. The cells come in close contact and move away from one another. (C) Brightfield image of a cell pair being tracked for the strain dynamics analysis in D. Cells are marked with white dotted lines and represented as C1 and C2. The red, green and yellow circles mark the beads being tracked for measuring the strain between C1 and C2, rear-end strain of C2, and rear-end strain of C1, respectively. (D) Strain as a function of time between C1 and C2 (black), rear-end of C1 (red) and C2 (blue). Strains are small (< 5%) even when the cells are very close to one another. The two red vertical dotted lines represent the time-window when the cells are closest to each other. When C1 exhibits 20% strain at its rear end, C2 exerts low strain on the matrix - evidence of a lack of interaction. (E) The distance between the cells decreases for a brief period (red vertical dotted lines), after which the cells move away from one another. Matrix strains between the cells were low irrespective of the distance. (F) SHG image between 2 such cells shows no collagen fiber bundle, implying no cell-cell interaction.

To determine whether such transient encounters induce matrix strain stiffening, we tracked embedded 2-µm beads in the collagen. Figure 1C shows a representative cell pair (C1 and C2), together with the strain measured between the cells and on their rear sides (Fig. 1D). The tensile strain between the cells remained low, reaching only ∼5%, comparable to that generated by isolated single cells(17) and substantially lower than that observed between interacting cells in high-density cultures. Moreover, the strain did not increase as the cells approached one another but fluctuated stochastically throughout the observation period (Fig. 1D), indicating that the intervening matrix did not undergo progressive strain stiffening and that the cells failed to enter the positive mechanical feedback loop.

Consistent with these measurements, second-harmonic generation imaging revealed no detectable collagen fiber alignment or densification between the cells (Fig. 1H), in sharp contrast to high-density cultures (Fig. 2). Thus, two isolated nearby cells are insufficient to establish persistent mechanical communication or remodel the surrounding matrix.

**Fig 2.**
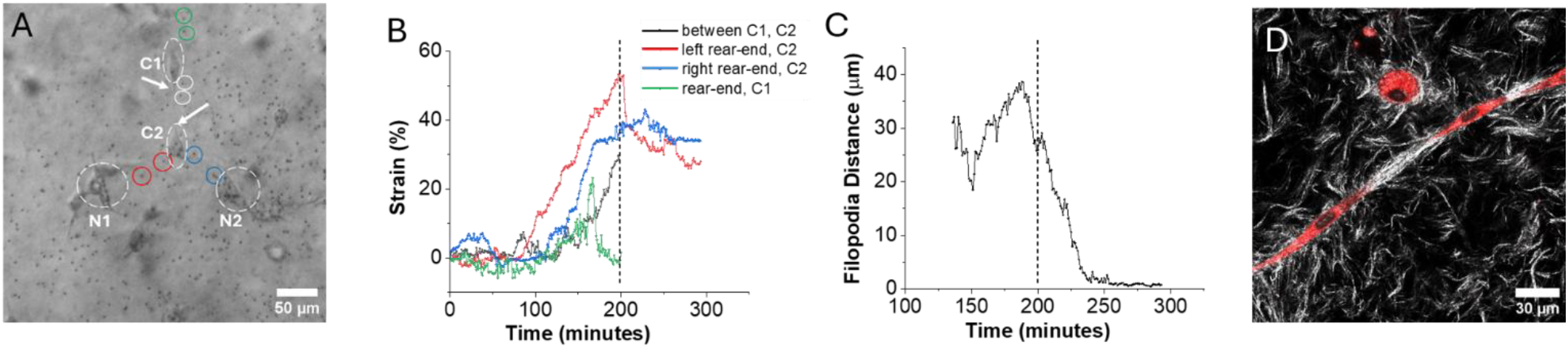
Strain dynamics between 3-cells show strong mechanical interaction leading to network formation in high-cell density collagen disk. (A) Brightfield image of the cells (C1, C2) being tracked over time for strain analysis in B. The white, red, blue and green circles mark the bead pairs tracked for measuring strain between C1 and C2, left and right rear-end strains of C2, and rear-end strain of C1, respectively. (B) Corresponding strains as a function of time between C1 and C2 (black), at left (red) and right (blue) rear end of C2, and at rear-end of C1 (green). In four cases, the strains increase sharply and reach values of 20-50%. (C) Distance between the filopodial tips of C1 and C2 as a function of time. When the cells are increasing matrix strain (B), there is a negligible change in the distance. At the 200th minute, strain between C1 and C2 reaches a peak, after which filopodia rapidly approach each other (tip to tip distance drops). The tips meet each other to form part of a network. (D) Images of two cells (CellTracker Red) at high cell density cell-collagen matrix participating in network formation. Confocal reflectance image of collagen shows fiber bundling between the cells.

### Opposing mechanical anchors enable neighboring cells to strain-stiffen the matrix and form stable networks

At high cell density, fibroblasts embedded in collagen-1 (2 mg/mL) initially migrate randomly but rapidly transition to persistent interactions, forming stable multicellular networks that compact the gel (17)(SI Movie 3). To test whether neighboring cells act as mechanical anchors, we quantified local ECM strains around interacting cell pairs using embedded 2-µm tracer beads (SI Movie 4).

Figure 2A shows two interacting cells (C1 and C2), each supported by nearby neighbors, such as cells N1 and N2 for C2, that provide opposing mechanical anchorage. Strain mapping revealed a progressive increase in tensile strain between C1 and C2, reaching ∼30% (Fig. 2B), well within the nonlinear strain-stiffening regime of collagen(33). At the same time, both cells generated similarly large strains on their rear sides, where neighboring cells acted as mechanical anchors. For example, C2 strained the matrix by ∼50% and ∼40% on its two rear sides, while C1 generated ∼25% strain on its rear side. Thus, each cell simultaneously strained the matrix on both sides.

This bilateral strain escalation coincided with persistent filopodial extension toward one another (Fig. 2C). During the first ∼200 min, the strain increased steadily while filopodia fluctuated about their initial separation. Thereafter, the filopodia rapidly converged, established stable contact, and remained connected until large-scale network formation prevented further imaging. Second-harmonic generation imaging of similarly staged samples revealed pronounced collagen fiber alignment and densification along the interaction axis (Fig. 2D).

Unlike isolated cell pairs, where strains remained weak and stochastic, neighboring cells in dense cultures contracted synchronously while supported by opposing mechanical anchors. This bilateral strain stiffening enabled persistent mechanical coupling and the formation of stable multicellular networks.

### Compression-induced collagen buckling drives gel compaction

Our hypothesis predicts that once a contractile cellular network forms, it compresses the collagen matrix enclosed within it. When the compressive stress exceeds the buckling threshold of collagen fibers, the matrix collapses, expelling interstitial fluid and producing macroscopic gel compaction.

To test this prediction, we performed time-lapse confocal reflectance microscopy of collagen fibers surrounding live cells under three conditions: (1) an isolated cell in a low-density gel, (2) a cell pair during the early stages of network formation in a high-density gel, and (3) multiple interacting cells in a compacting high-density gel (Fig. 3A–F). Fibers remained relatively straight around isolated cells, resembling those in cell-free collagen, but became increasingly buckled as cellular networks formed and the gel compacted.

**Fig 3.**
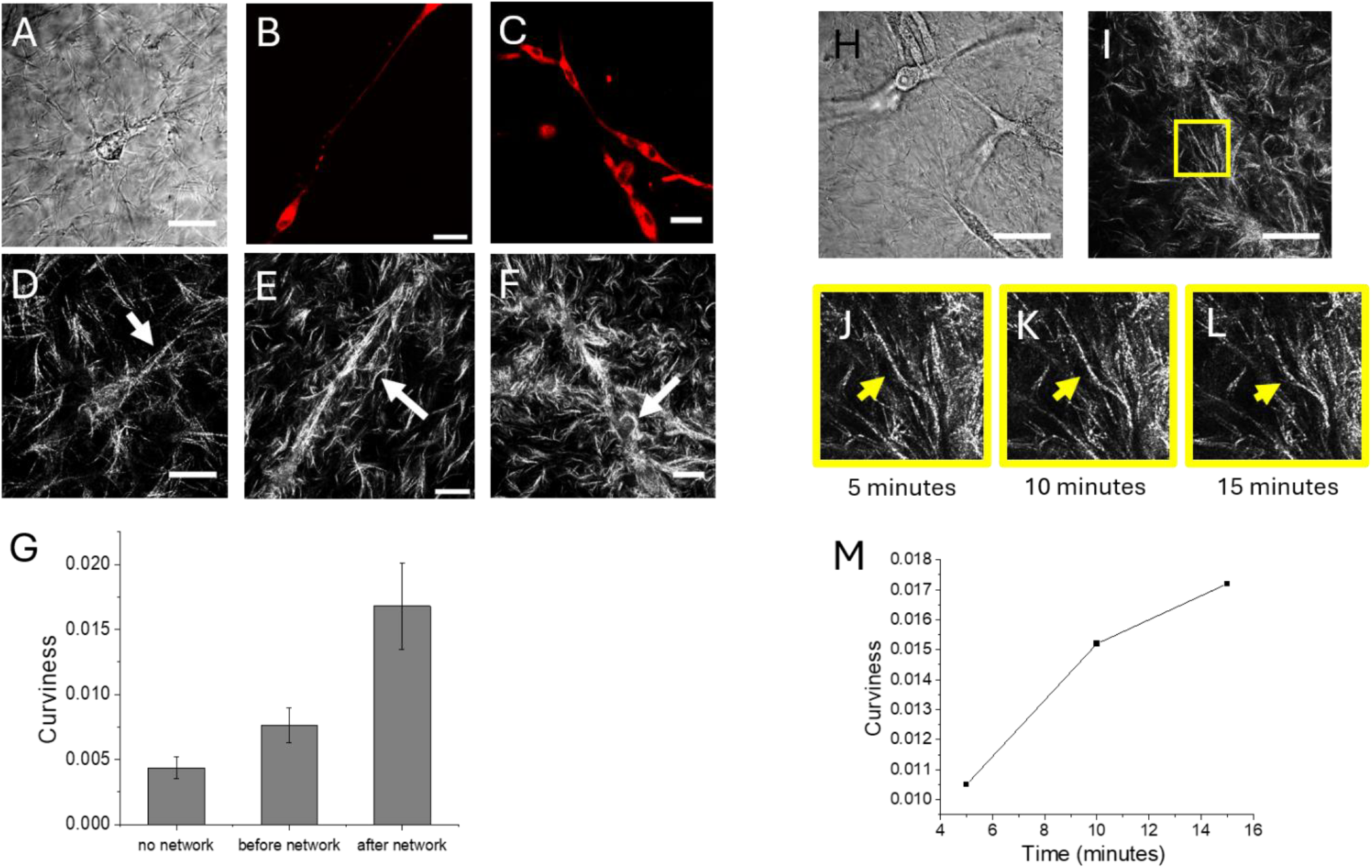
Buckling of collagen fibers leads to compaction of collagen disks with fibroblasts. (A-C) brightfield and confocal images of a single cell (A), 2 cells at early stages of network formation (B), multiple cells at later stages of network formation (C). (D-F) Corresponding collagen fiber images from confocal reflectance microscopy, scale bar 30 µm. (G) Average curviness of collagen fibers for cases described in D-F. (H-I) Brightfield and confocal reflectance images of the region of the collagen disk with live cells being imaged over time for tracking the fibers marked in the region with a yellow rectangle in I. (J-L) Confocal reflectance images of the fiber being tracked at different time points. (M) Curviness of the fiber as a function of time.

We quantified fiber buckling by measuring the **curviness (Methods)**, 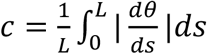, where *L* is the length of a fiber, *s* is the coordinate along the fiber, *θ*(*s*), is the angle at *s*. Mean curviness increased from ∼0.005 rad/µm around isolated cells to ∼0.017 rad/µm in compacting cellular networks (Fig. 3G), indicating a threefold increase in fiber buckling.

We next visualized buckling directly by tracking individual collagen fibers adjacent to interacting cells during network formation (Fig. 3H–M). Fiber curviness increased from ∼0.010 to ∼0.017 rad/µm within 15 min. Together, these observations demonstrate that macroscopic collagen compaction is driven by compression-induced fiber buckling following the formation of a contractile multicellular network(34–37).

### Persistent filopodial extension is guided by opposing mechanical anchors

We next asked whether persistent filopodial extension is directed by mechanical cues alone. To isolate the mechanical contribution of neighboring cells, we replaced them with rigid, chemically inert polystyrene (PS) beads (15 µm diameter) coated with collagen (Methods). The beads adhered to the ECM but remained biologically inert, thereby serving as localized mechanical anchors. Local ECM strains were quantified by tracking embedded 2-µm tracer beads (Fig. S3).

A cell positioned between two opposing beads generated >10% tensile strain toward both beads simultaneously, provided that each bead was within ∼100 µm (Fig. S3A–C). Under these conditions, the cell polarized along the bead axis and extended a persistent filopodium toward one of the beads (Fig. S3C). In contrast, when a bead was present on only one side, the strain toward the bead reached ∼15% transiently, whereas the strain on the opposite side remained low (2–3%) (Fig. S3D,E). The cell exhibited only transient cycles of filopodial extension and retraction, with no persistent advance toward the bead (Fig. S3F).

These results demonstrate that opposing mechanical anchorage, rather than on one side alone, is sufficient to trigger persistent mechanosensing and directed filopodial extension in the absence of biochemical signaling.

### A critical-force theory of collective cell–ECM interactions

The experiments suggest a simple physical picture. Mechanosensitive cells embedded in a soft fibrous matrix pull on and release collagen fibers while migrating randomly. In the absence of opposing mechanical anchors, matrix deformation remains localized and transient, preventing sustained strain stiffening. However, when a cell is supported by neighboring cells or rigid objects on both sides, it can strain-stiffen the matrix bilaterally, increasing the effective stiffness that it senses. This promotes cell polarization, enhanced contractility, and further matrix stiffening, establishing a positive force–strain–force feedback loop. Neighboring cells can likewise enter this feedback loop if they are themselves supported by opposing anchors, allowing a contractile multicellular network to emerge. Tensile forces and matrix stiffness then percolate through the network, compressing the enclosed matrix until collagen fibers buckle, producing macroscopic gel compaction. Thus, although two neighboring cells are insufficient for persistent mechanical interaction, three mechanically coupled cells can form the minimal mechanically stable unit capable of sustaining collective contraction.

These observations raise a fundamental question: **what initiates the transition from stochastic probing to persistent force generation?** We propose that this transition is governed by a **critical force**, *F*_*cr*_. When cell force, *F*_*c*_ < *F*_*cr*_, cells repeatedly assemble and release transient adhesions, producing fluctuating forces. If matrix stiffness enables the contractile force to exceed *F*_*cr*_, adhesions stabilize and the cell increases its force continuously, reaching a matrix-dependent stall force, *F*_*c*_ = *F*_*stall*_. The cell enters a persistent contractile state. Bilateral mechanical anchorage promotes this transition by supporting the increase in matrix strain and hence strain stiffening. The cell senses the increase in the effective matrix, further increasing its force in a positive feedback loop.

In our proposition, **critical force** is an intrinsic mechanical property of a cell type. *F*_*cr*_ defines the threshold force required for a cell to transition from stochastic probing to persistent force generation. Different cell types are therefore expected to possess different critical forces, reflecting differences in their mechanosensitive machinery and adhesive dynamics. Whereas the effective stiffness depends on the extracellular environment that determines the cell stall force. Collective behavior emerges only when the effective stiffness created by the surrounding matrix and neighboring cells enables the contractile force to exceed this intrinsic threshold. Thus, the onset of collective remodeling is governed by the interplay between an intrinsic cellular property, the critical force, and an extrinsic environmental property, the effective matrix stiffness. Theoretically, if *F*_*stall*_ ≫ *F*_*cr*_, the cell is expected to transition from stochastic probing to a persistent contractile state. The reverse is expected if *F*_*stall*_ ≪ *F*_*cr*_.

This hypothesis cannot be explored experimentally over the broad parameter space defined by matrix stiffness, cell density, and mechanical boundary conditions. We therefore develop a minimal one-dimensional theory that incorporates nonlinear ECM mechanics and the critical-force hypothesis to predict collective cell–ECM interactions across diverse conditions. The theory yields experimentally testable predictions, which we validate in the following sections.

### Theoretical model of cell–ECM interaction

Our theory is rooted in the concept that collective cell–ECM interactions are governed by two physical principles: **an intrinsic critical force**, which defines the mechanical identity of a cell, and an **effective matrix stiffness**, which characterizes the mechanical environment experienced by the cell. We developed a minimal one-dimensional theory based on these principles to predict collective behavior over a broad range of matrix stiffnesses, cell densities, and boundary conditions.

### Critical-force theory

A living cell adheres to surrounding ECM fibers and actively contracts. Fibers aligned with the direction of contraction are placed under tension, whereas the surrounding matrix experiences compression. The cell forms a local mechanical zone of influence (Fig. 4A).

**Fig 4.**
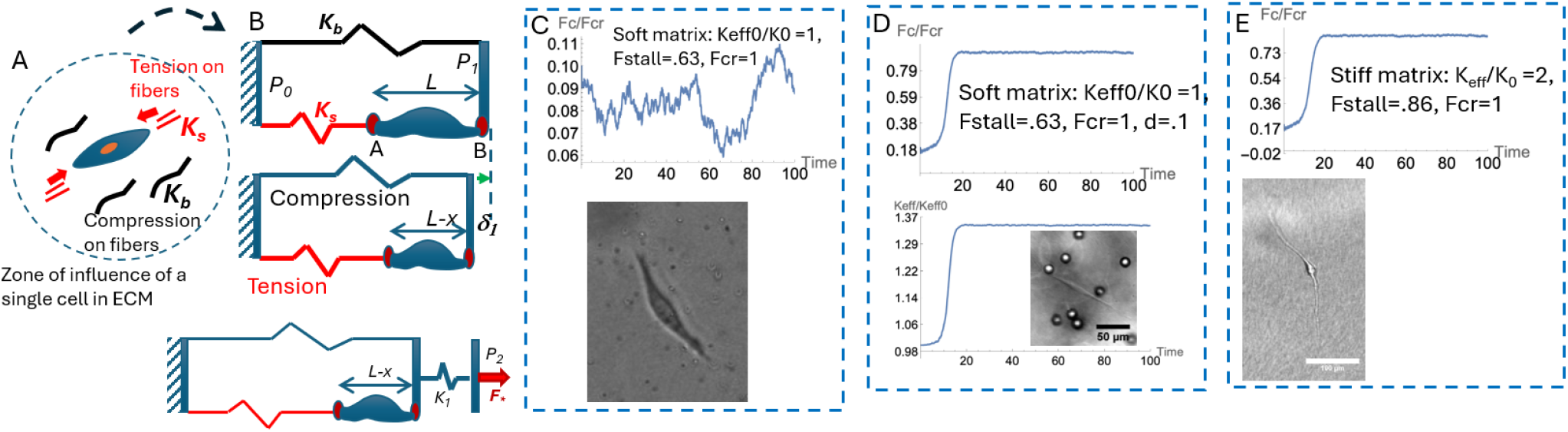
Minimalistic model of a cell-ECM unit, model predictions, and experimental verifications. (A) A single cell embedded in a fibrous ECM applies a force dipole. (B) 1-D model of the cell-ECM system, referred to as the CE unit. It consists of 2 rigid plates, *P*_*0*_ and *P*_*1*_, as its boundaries, a tensile and a compressive spring mimicking the tensile and compressive resistance of the ECM, and a cell. It is connected to a 3^rd^ plate, *P*_*2*_, through a non-linear spring *K*_*1*_ to represent strain stiffening and to account for its interaction with neighbors. The cell applies a contraction, x(t), and shortens its length from its rest length, L. The cell cannot stretch. It experiences a tensile force, Fc, referred to here as the cell force. (C) The model predicts stochastic cell force in a soft matrix (top). Experimental image of a fibroblast in soft 2 mg/ml collagen (below). (D) The model predicts that the same cell in the soft matrix, but close to rigid boundaries on both sides, transitions from stochastic to deterministic stall force mediated by bilateral strain stiffening. Inset shows an experimental fibroblast polarizes and becomes stable between rigid beads in 2 mg/ml collagen. (E) The model predicts that the same cell as in (C) and (D) in a stiffer matrix transitions from stochastic to deterministic stall force without any further stiffening of the matrix (top). Experiment shows a single fibroblast becomes highly polarized in stiffer (8 mg/ml) collagen.

We represent the cell and the ECM by a cell–ECM (CE) unit, a mechanical analogue. The cell is represented by an active contractile actuator, while the surrounding ECM by two **linear elastic** elements: a tensile spring (*K*_*s*_) representing the collagen fibers under tension and a compressive spring (*K*_*b*_) representing the remaining surrounding matrix (Fig. 4B). Although simplified, this representation captures the local mechanics of an isolated cell while deliberately separating it from nonlinear multicellular interactions.

The actuator shortens by an amount *x*(*t*), producing tensile and compressive forces in the springs. Corresponding cell force, *F*_*c*_(*x*), can be obtained from force balance (Methods). The effective stiffness sensed by the cell can be derived from 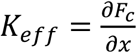. We model cell stall force as a function of *K*_*eff*_, as 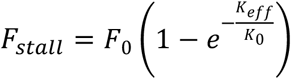. For the isolated CE unit, *K*_*eff*_ is only a function of *K*_*s*_ and *K*_*b*_, and the unit behaves as a linear elastic system.

### Nonlinear coupling

Collective behavior is introduced by coupling neighboring CE units through nonlinear tensile springs representing the ECM fibers spanning adjacent cells or nearby rigid anchors. These springs strain-stiffens as *F*_∗_ = *F*_0_(Δ/*d*)^3^, where *F*_∗_ is the spring force, Δ and *d* are its stretch and length respectively, *F*_0_ is a force constant. Note that, *d* defines the proximity between the cell and its neighbor.

Coupling increases the effective stiffness, *K*_*eff*_, experienced by the cell. The degree of increase depends on the activities of the cell and its neighbor (their contractile states, *x*(*t*)), as well as the distance between them. Hence, even if the neighbor is an inert rigid object, it can support the stiffening due to the contractility of the cell. Coupling vanishes when *d* → ∞. *K*_*eff*_ can be computed from force balance and compatibility (Methods).

The critical-force hypothesis introduced above is implemented phenomenologically by assigning each cell two quantities: intrinsic critical force, *F*_*cr*_, and a matrix-dependent extrinsic stall force, *F*_*stall*_(*K*_*eff*_). Cell activity depends on these two quantities. We model the activity through its contraction dynamics, *x*(*t*), consisting of a stochastic and a deterministic component:

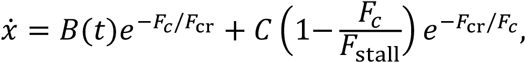

where *B*(*t*)represents stochastic fluctuation and C is a constant. Below the critical force, stochastic fluctuations dominate. Above the critical force, deterministic contraction dominates and the cell approaches its stall force.

### Theoretical predictions and experimental validations

#### Single cell–ECM (CE) model

##### Soft matrices support stochastic force generation

In a soft matrix, an isolated cell experiences a low effective stiffness and therefore may not generate sufficient force to exceed its intrinsic critical force, *F*_*cr*_. Then, *F*_*cr*_ ≫ *F*_*stall*_. The model predicts that the cell remains in a stochastic probing state, producing fluctuating forces and weak, transient matrix deformations without appreciable strain stiffening (Fig. 4C). This prediction agrees with experiments showing that NIH 3T3 fibroblasts embedded in soft collagen (2 mg/mL, 0.5 kPa) generate stochastic matrix strains while migrating randomly(17). Similar is the case in soft linear elastic ColT gels (Fig. S2).

##### Mechanical boundary conditions can substitute for matrix stiffness

The model predicts that an isolated cell in a soft matrix can transition to persistent contraction if rigid mechanical anchors are present on opposite sides. Bilateral anchorage allows the cell to increase the effective stiffness of the matrix, escalate its contractile force in a feedback loop process to exceed the critical force and reach the persistent contractile state. Experiments confirmed this prediction: cells positioned between two collagen-coated rigid beads generated large bilateral matrix strains, polarized along the bead axis, and extended persistent filopodia toward the anchors (Fig. 4D; Fig. S3A, B). In contrast, a single nearby anchor failed to induce persistent polarization.

##### Increasing the effective stiffness stabilizes force generation

The model predicts that for the same cell type as above (same *F*_*cr*_), if effective stiffness is sufficiently large so that the corresponding stall force greatly exceeds the critical force, *F*_*cr*_ ≪ *F*_*stall*_, then the cell transitions from a stochastic to a persistent state, approaching the stall force irrespective of whether the stiffness originates from the matrix itself or from mechanical constraints. This prediction is confirmed experimentally. NIH 3T3 fibroblasts embedded in stiff collagen (8 mg/mL, *E*∼5 *KPa*) became largely stationary, remained polarized for more than 14 h, and exhibited only small filopodial fluctuations about a fixed axis (Fig. 4E). Similar behavior was observed in stiff (10 kPa) linear elastic ColT gels.

### Minimal network model

To investigate collective behavior, we connect three CE units through nonlinear strain-stiffening springs (Fig. 5A). Three cells constitute the smallest mechanically connected network in which every cell is supported by opposing mechanical anchors. The matrix enclosed within a living cellular network is represented by a compressive spring exhibiting elastic, buckling, and post-buckling stiffening behavior (Fig. 5B). Compression of this spring provides a measure of gel compaction. The mathematical details are provided in Methods.

**Fig 5.**
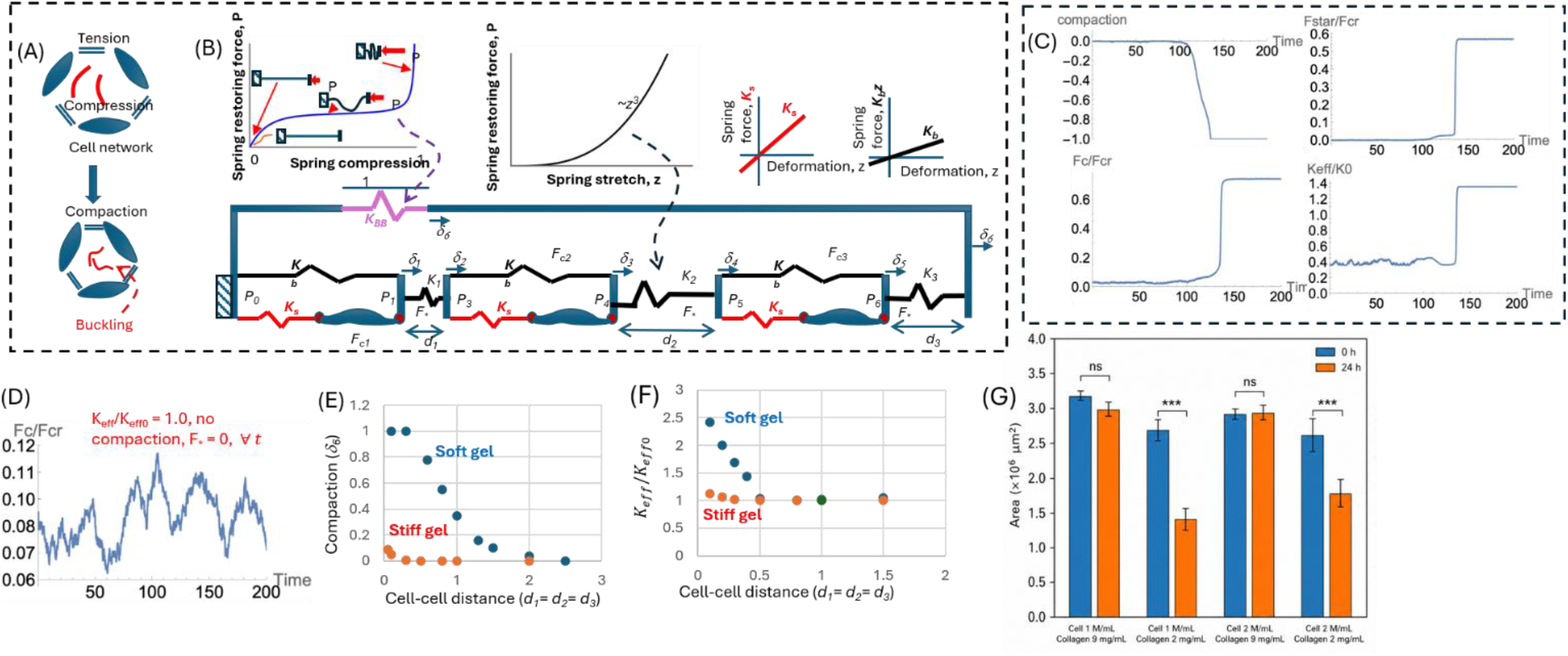
Minimalistic model of a cell-ECM system with 3 cells, model predictions, and experimental verification. (A) Schematic of a 3-cell network. (B) 1-D model with 3 cell-ECM units connected by non-linear springs, *K*_*1*_, *K*_*2*_, *K*_*3*_, allowing strain stiffening. The non-linear spring *K*_*BB*_ represents ECM under compression. *K*_*BB*_ has a unit length. Its compression, *d*_*6*_, represents gel compaction; (C) Model predictions on gel compaction, cell force, cell-cell interaction force, and matrix effective stiffness with time when cells are nearby (*d*_*1*_ *= d*_*2*_ *= d*_*3*_ *= 0*.*1*). Cells transition from stochastic to steady stall force; (D) If one of the 3 cells is far from the other two (*d*_*1*_ *= 0*.*1, d*_*2*_ *= 100, d*_*3*_ *= 0*.*1*), then cells fail to interact, and their forces remain stochastic. (E-F) Model predictions on compaction and ECM stiffening with cell-cell distance. Compaction and stiffening are significantly higher in soft gel compared to those in stiff gel. (G) Experimental validation of model predictions in (E). Fibroblasts (2 M/ml) in free-floating gel discs compact soft gel (2 mg/ml) but not stiff gel (8 mg/ml).

The three-cell model predicts that collective matrix remodeling emerges only when cells are sufficiently close to establish opposing mechanical anchorage. Fig. 5C shows the theoretical prediction of the evolution of cell force, effective matrix stiffness, cell–cell interaction force, and gel compaction as the cell–cell spacing (*d*_1_ = *d*_2_ = *d*_3_) is varied. At high cell density (small *d*_*i*_), cells initially generate stochastic forces but rapidly strain-stiffen the ECM, increase the effective stiffness, and exceed the critical force, all of which are facilitated by cell-cell interactions and the positive feedback loop. The cells subsequently approach their stall force. Their tension is balanced by the compression spring, mimicking ECM under compression. The spring buckles allowing large compaction. The spring then stiffens sharply under compression, resisting any further compaction. As *d*_*i*_ increases, both strain stiffening and compaction decrease, disappearing beyond a critical separation (Fig. 5E,F). These predictions qualitatively reproduce experimental observations on collagen compaction with cell density (17).

The model further predicts that **two neighboring cells are insufficient** to initiate collective contraction. When one of the three cells is moved far away, *d*_2_ ≫ *d*_1_, *d*_3_, bilateral mechanical anchorage is lost. Consequently, cell force remains stochastic, while the effective stiffness remains unchanged (i.e., no strain stiffening), the interaction force vanishes, and gel compaction does not occur (Fig. 5D). These predictions agree with experiments showing that isolated cell pairs generate only weak, stochastic matrix strains and ultimately separate without establishing stable mechanical interactions (Fig. 1).

Finally, the model reveals that **high cell density alone is not sufficient for collective behavior**. Although a stiff matrix promotes persistent contraction of individual cells, it suppresses the additional strain stiffening required for neighboring cells to increase one another’s effective stiffness. As a result, cells remain mechanically isolated despite being closely packed, preventing the positive force–strain–force feedback loop from propagating through the network. Consequently, the model predicts little cell–cell interaction and negligible gel compaction in stiff matrices even at high cell density (Fig. 5E). Experiments confirmed this prediction: NIH 3T3 fibroblasts (2 M/mL) compacted soft collagen gels (2 mg/mL) by approximately 50% within 24 hrs but failed to compact stiffer collagen gels (9 mg/mL) over the same period (Fig. 5G).

## Discussion

This work advances the evolving understanding of cell–ECM interactions. Collagen is the dominant extracellular matrix (ECM) in many tissues, where reciprocal cell–ECM crosstalk regulates tissue morphogenesis, wound healing, and skin grafting (14). Collagen stiffens under strain through fiber alignment rather than increased crosslinking (25, 38). In vitro, fibroblasts compact collagen gels by more than 80% without chemical degradation when cell density exceeds a threshold (13, 16, 17). Individual cells can locally strain-stiffen collagen, increasing matrix stiffness, further promoting contractility in a positive feedback loop process triggering cell polarization (19, 22). These observations raise a central question: what initiates this positive feedback between cells and the ECM? Is there a unifying principle of cell-ECM interaction?

Our results suggest that cell–ECM interactions are governed by the competition between two physical parameters: intrinsic critical force, *F*_*cr*_, which defines the mechanical identity of a cell, and the effective matrix stiffness, which determines cell’s stall force, *F*_*stall*_. When the effective stiffness is sufficient for the contractile force *F*_*c*_ to exceed *F*_*cr*_, cells transition from stochastic probing to persistent force generation. This transition can occur either in intrinsically stiff matrices or in compliant fibrous matrices that cells strain-stiffen through their own contractility.

In high-cell-density matrices, neighboring cells amplify this process by providing opposing mechanical anchorage that enables bilateral strain stiffening. Pairwise proximity alone cannot support this stiffening. Collective behavior emerges only when cells become mechanically connected through the ECM, increasing the effective stiffness experienced by neighboring cells. This mechanical reinforcement promotes filopodial extension toward neighboring cells, enables percolation of long-range tensile force, and drives the transition from cell’s stochastic probing to persistent force generation. Once a tensile network is established, collagen buckling and post-buckling stiffening produce macroscopic gel compaction (Fig. 6).

**Fig 6.**
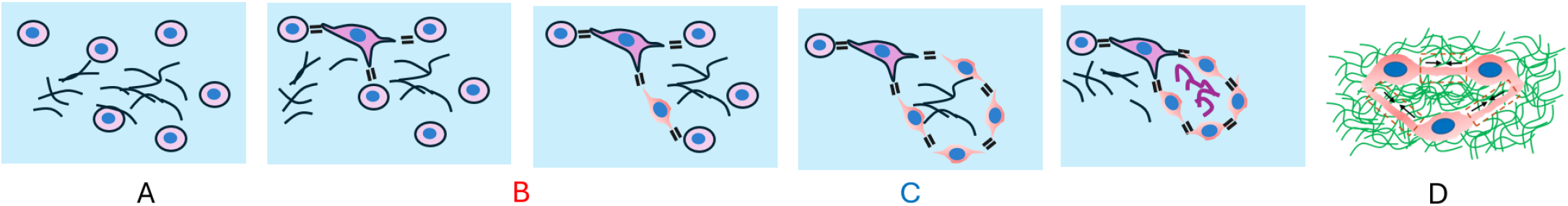
Conceptual model of cell-ECM interaction and gel compaction at high cell density. Each cell serves as a mechanical anchor for its neighbor. Initially (state A), cells generate a stochastic force on the ECM. One of the cells pulls the matrix using 3 of its neighbors, strain-stiffening the matrix in between, resulting in an increase in cell force. The cell enters the positive feedback loop (state B). The neighbors, being active, begin to polarize towards each other, further increasing the force and forming a network or a loop through which tension percolates. The emergence of this network is the first phase transition (state C). ECM trapped between the loop is compressed, balancing the tension. If the compression exceeds a threshold, ECM fibers buckle, and gel volume decreases rapidly (state D). (E) At least 3 cells are needed to form the cell network.

This framework shifts mechanosensing from a purely local material property to a network property. It identifies mechanically connected topology as the missing architectural requirement for long-range collective remodeling and explains why compliant strain-stiffening matrices can generate stronger collective behavior than uniformly stiff matrices.

Despite its simplicity, the theory explains most of the observations on cell-ECM interactions reported to date. It describes why three mechanically coupled cells form the smallest stable contractile unit, why rigid inclusions promote gel compaction by acting as mechanical anchors, and why gels spanning orders of magnitude in cell density compact to similar final volumes once a mechanically connected network is established. More broadly, it predicts that collective remodeling is governed jointly by cell type, matrix architecture, cell density, and mechanical boundary conditions through the competition between intrinsic critical force and cell stall force determined by network-generated effective stiffness.

These principles likely extend beyond collagen gel compaction to tissue morphogenesis, wound healing, fibrosis, and tumor progression. In each case, tissue-scale remodeling emerges when mechanically connected cell–ECM networks generate sufficient effective stiffness for cells to overcome their intrinsic critical force, providing a unified physical framework linking local cellular forces to tissue-scale mechanics.

## Methods

### Cell culture

We used NIH 3T3 fibroblasts(passage number 5-10) for our experiments. Cells were cultured in flasks in media consisting of 89% v/v Dulbecco’s Modified Eagle’s Medium (DMEM) (Corning® Inc., Corning, NY), 10% v/v Fetal Bovine Serum (FBS) (ThermoFisher Scientific™, Waltham, MA) and 1X Penicillin Streptomycin (Lonza®, Basel, Switzerland) until reaching 70-80% confluency. Cells were detached from the flasks using 0.05% Trypsin-EDTA (Gibco™, Waltham, MA) and centrifuged at 150 g for 5 minutes to obtain a cell pellet. The cells were next resuspended in cell culture media and counted using a hemocytometer (Hausser Scientific™). Next, cell suspension of double the desired cell density was prepared with cell culture media.

### Fibrous collagen disc preparation

Type I collagen with a density of 4 mg/ml (Corning® Inc., Corning, NY) was prepared from a stock solution according to the manufacturer’s protocol. Briefly, collagen was diluted with ice-cold deionized water and 10X phosphate buffer saline (PBS) (Lonza®, Basel, Switzerland). The solution was neutralized by adding 10N NaOH (Sigma Aldrich, Saint Louis, MO) to achieve a pH of 7.2-7.4. Equal volumes of collagen solution and cell suspension containing double the desired cell densities were mixed on ice to achieve a cell-collagen mixture of the desired density, with a final collagen concentration of 2 mg/ml. The cell-collagen mixture was dispensed inside a PDMS scaffold containing cylindrical holes with a diameter of 2mm and a thickness of 0.5mm. The cell-collagen mixture forms a film inside the holes, which was then incubated at 37 °C for 25 minutes until the collagen solution polymerized, forming disc-shaped gels. After polymerization, cell-culture media were poured on top of the discs to detach them from the Pluronic-coated scaffolds. The scaffolds were removed, and free-floating collagen discs were obtained.

### Linear elastic Col-T gel preparation

To prepare the linear elastic nonfibrous 3D matrix with cells, we used Col-T gel (101 Bio) which is a commercially available linear elastic hydrogel with tunable elastic properties. It consists of 2 components – a matrix component A and a transglutaminase-based crosslinking component B. They were mixed in different ratios according to the manufacturer’s protocol to achieve a bulk elastic modulus of 0.5 kPa and 10kPa after the gels polymerized. 3T3 cells were mixed with the liquid Col-T gel mixture in a similar method to collagen. To improve cell spreading and attachment, 0.7 mg/ml fibrinogen (diluted in PBS) was added to the Col-T gel mixture, which has been previously shown to promote cell attachment in 3D linear elastic polyethylene glycol (PEG) gels(39). The mixture was then dispensed inside the PDMS molds and incubated at 37 °C for 1 hour. The gels were then detached from the scaffold using a needle to obtain the free-floating 0.5 kPa and 10 kPa Col-T gel discs.

### PS-bead preparation

Inert polystyrene (PS) beads (Bangs Laboratories, Fishers, IN) of 15 µm diameter were coated with collagen before mixing them with cell-collagen mixture according to manufacturer’s protocol. Briefly, the desired volume of bead stock solution was centrifuged at 200g for 5 minutes to obtain a bead pallet with desired number of beads. After aspirating the liquid suspension, the bead pallet was re-suspended in 0.1% collagen solution in PBS for 20 minutes. This solution was then centrifuged again, and the solvent was aspirated to obtain collagen-coated PS beads. For the experiments with different cell densities, a bead density of 1 million/ml of collagen was used.

### Live cell confocal reflectance microscopy

High-resolution confocal reflectance time-lapse imaging was done on live collagen samples to visualize collagen fibers adjacent to cells protruding towards each other in a compacting disc over time. This was done using a 40x water immersion objective and a 633 nm laser in a laser scanning confocal microscope (LSM 880, Zeiss) equipped with an environment control chamber for cells. Live cells were imaged by labeling them with a red fluorescent dye (Cell Tacker, Invitrogen, excitation 561 nm). Images were taken every 5 minutes for 8 hours.

### Measurement of curviness of buckled collagen fibers

The buckling of the collagen fibers from the confocal reflectance images were quantified by measuring the curviness of the fibers from the images. We define curviness, *c*, of a fiber AB of length L as,

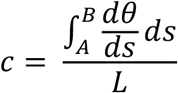

Where θ is the angle of the tangent at arc length, s, from the end A of the fiber. A threshold was applied on the confocal images of the collagen fibers using image processing software Fiji before feeding them in a custom MATLAB script that can segment the fibers in the images and measure the curviness of each fiber by applying the above equation. The script calculates the average curviness of the region of interest and gives the value as the output.

### Image and data acquisition

All the brightfield time-lapse images of cells were taken with a Olympus IX81 widefield microscope using a 10x objective equipped with an environment-control chamber for cells. The tracer bead displacements for strain analysis were tracked using a software called Tracker, which tracks the coordinates of the beads as a function of time by applying a template-matching algorithm.

### Theoretical model of cell-ECM interaction

#### Cell-ECM unit

Each cell-ECM (CE unit) unit represents a single cell in ECM. It consists of an actuator AB, mimicking the living cell, and two linear springs with spring constants *K*_*s*_ and *K*_*b*_ (Fig. 4B). The springs are termed as *K*_*s*_ and *K*_*b*_. The CE unit has 2 rigid plates, *P*_0_ and *P*_1_, defining its left and right boundaries respectively. *P*_0_ serves as a reference and hence is fixed in location. The cell applies a contractile force dipole, shortening its length by *x*(*t*) (Fig 4B). It can only contract, not stretch from its rest length. Hence, *x*(*t*) ≥ 0 for ∀*t*. Contraction *x*(*t*) induces a stretch in *K*_*s*_ and compression in *K*_*b*_, resulting in a tensile force, 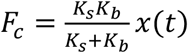 on the cell AB, and a displacement of *P*_1_ towards *P* by 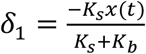. The cell “feels” an effective stiffness, *K*_*eff*_, while contracting by *x*(*t*), given by 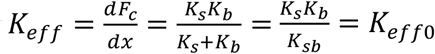, where *K*_*sb*_ = *K*_*s*_+ *K*_*b*_. Here *K*_*eff*0_ represents the intrinsic linear elastic stiffness of the matrix.

In order to account for an external force on CE due to a nearby cell or a boundary, consider an external force, F*, applied on *P*_1_ towards right. *F*_∗_ is always tensile, due to contractile nature of cells. Then force balance and displacement compatibility requires:

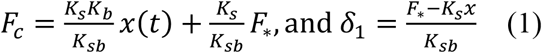

For fixed 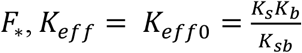 is independent of *F*, i.e., implying no cell-neighbor interaction.

To account for cell-neighbor interactions involving non-linear ECM stiffening, consider a rigid plate, *P*_2_, connected to *P*_1_ by a non-linear spring, *K*_1_ (Fig. 4D). Later, we will replace *P*_2_ with a cell-ECM unit. Let *d*_1_ be the initial gap between *P*_1_ and *P*_2_. Let *δ*_2_ be the displacement applied to *P*_2_. Recall that *P*_0_ is held fixed. Then the spring, *K*_1_, is stretched by (*δ*_2_ − *δ*_1_), where *δ*_1_ is the displacement of *P*_1_ due to the combined effect of cell contraction *x*(*t*) and applied *δ*_2_. We model the restoring force of the non-linear spring, *K*_1_, as

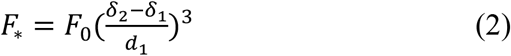

where *F*_0_ is a constant. Force balance requires:

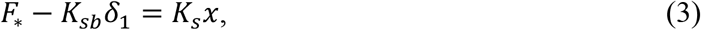

### Effective stiffness

One can solve *δ*_1_(*x*) from Eqs 2 and 3 for a given *δ*_2_. In response to applied *δ*_2_, both *K*_*eff*_ and *F*_*c*_ increase mimicking the interaction between a living cell and its neighbor:

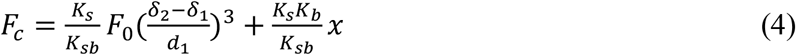

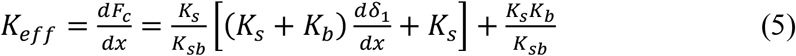

Eqs. 4 and 5 allow to predict possible response of a living cell in a wide range of ECM stiffness and its interaction with neighbors.

### Critical-force dynamics

#### Stall force, *F*_*stall*_

mechanosensitive cells increase their contractile force after engaging with their substrates reaching steady force, *F*_*stall*_, depending on cell type and substrate stiffness, *K*_*eff*_ (40). We model this steady force as:

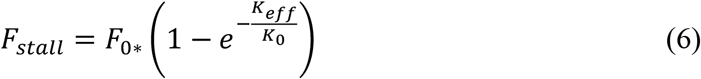

Here *F*_0∗_, *K*_0_ are constants.

#### Critical force, *F*_*cr*_

*F*_*cr*_ determines the stability of cell force dynamics. When *F*_*c*_ ≪ *F*_*cr*_, cell contraction 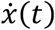 is stochastic. When *F*_*c*_ → *F*_*cr*_, cell force tends to become monotonic and deterministic. In this case, cell may continue to increase force to reach *F*_*stall*_, and 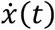 approaches zero. We model this stochastic probing to persistent force generation state by defining the rate of cell contraction, 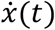 as follows:

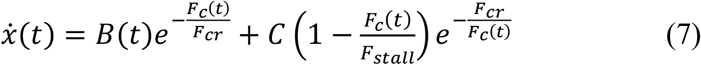

Here, *C* is a constant with the unit of velocity, and *B*(*t*) represents a stochastic function ensuring 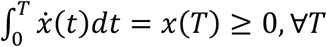.

### Minimal network model

To capture collective behavior of multi cellular systems in a reduced framework, we model the system as a one-dimensional arrangement of three CE units (Fig. 5A). The units are separated by distances *d*_*i*_, i = 1,2,3, connected by nonlinear springs *K*_*i*_, i = 1,2,3, of lengths *d*_*i*_ characterized by their force response, 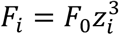, where *z*_*i*_ = (*stretch of spring i*/*d*_*i*_)^3^. For example, force in spring *K*_2_ is *F*_2_ = *F*_0_((*δ*_4_ − *δ*_3_)/*d*_2_)^3^ (Fig. 5B). Here, *δ*_*i*_, *i* = 1, … ,6, are the displacements shown in Fig. 5B. A compressive spring *K*_*BB*_ connects the first and the 3^rd^ units, closing the loop. Large *d*_*i*_ represents low cell density. Each cell contracts by an amount *x*_*i*_(*t*), *i* = 1,2,3, generating tensile force, *F*_∗_, in each of the springs *K*_1_, *K*_2_, and *K*_3_. *F*_∗_ is balanced by an equal compressive force on *K*_*BB*_ compressing it by *δ*_6_(< 0). *F*_∗_ gives a measure of cell-cell interaction. The spring *K*_*BB*_ is of length unity. Under small compression, it behaves linearly. With increasing compression, it buckles. With further compression, it stiffens rapidly as illustrated in Fig. 5B.

Force-displacement relation for *K*_*BB*_ is modeled as:

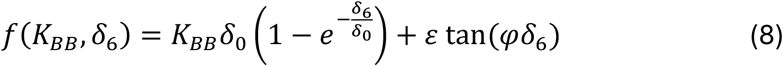

*δ*_0_ (< 0), *ε*, and 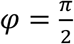 are constants. *δ*_6_ gives a measure of compaction in the 3-cell model.

We solve *F*_∗_(*t*) for given *d*_*i*_ and *x*_*i*_(*t*). Gel compaction begins when *F*_∗_(*t*) exceeds critical buckling force for *K*_*BB*_

To solve for *F*_∗_(*t*), we use force balance and displacement compatibility. Eq. 3 gives

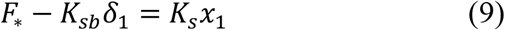

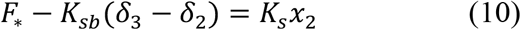

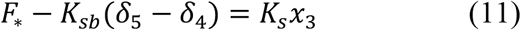

From Eq. 2 gives

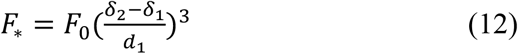

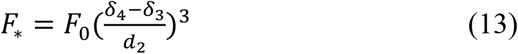

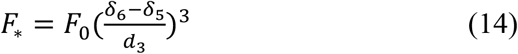

Eq. 8 gives

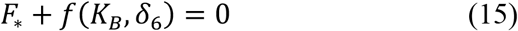

The 7 unknowns, i.e., *δ*_*i*_, *i*, … , 6, and *F*_∗_ can be solved numerically from the above 7 independent equations giving *δ*_*i*_(*t*) = *δ*_*i*_(*x*_*j*_(*t*), *d*_*j*_) and *F*_∗_(*t*) = *F*_∗_(*x*_*j*_(*t*), *d*_*j*_), *j* = 1,2,3. Cell forces for each cell-ECM unit can be determined from Eq. 4 as

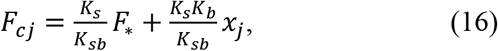

with corresponding effective stiffness,

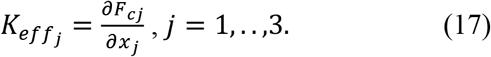

Stall force for each CE unit is then, from Eq. 6 is

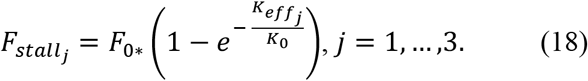

Finally, increments in *x*_*j*_(*t*) are given by Eq. 7

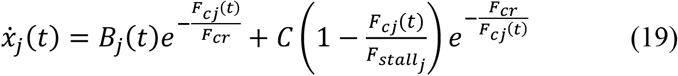

We apply the 3-cell model to explore a wide range of scenarios of cell-ECM interactions.

### Numerical implementation

Software used: **Mathematica**

Random noise generation, B(t) in Eq. 7:

We used Ornstein–Uhlenbeck (OU) type process. It describes a mean reverting random walk given by:

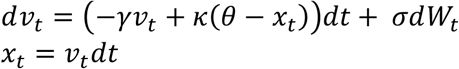

Where:

*x*_*t*_ is the stochastic process at time t

*κ*(*θ* − *x*_*t*_) is a mean-reverting force pulling *x*_*t*_ towards *θ* (=0.1), the long-term mean, *κ* (=0.5) is the speed of mean reversion.

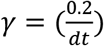 is velocity damping

*σdW*_*t*_ is Gaussian noise

*σ* (=.1) is magnitude of random fluctuations

Boundary condition applied so that *x*_*t*_ > 0.1.

*dt* is the time increment

### Model parameters used in the simulations

*K*_0_=1

*F*_0_=1

*ε* = .001

*φ* = *π*

*δ*_0_ = −.01

*F*_0∗_=1

Time step dt = 1/100

### Numerical implementation for single cell behavior

Fig. 4E1

*K*_*s*_ = 3, *K*_*b*_ = 1.5, *K*_*BB*_ = 0, *F*_*cr*_ = 1, *d*_1_ = 1000, *d*_2_ = 1000, *d*_3_ = 1000

*F*_*stall*_ = 0.63, *K*_*eff*0_ = 1, *K*_*eff*_ = 1, *δ*_6_ = 0.

Fig. 4F1

*K*_*s*_ = 6, *K*_*b*_ = 3, *K*_*BB*_ = 0, *F*_*cr*_ = 1, *d*_1_ = 1000, *d*_2_ = 1000, *d*_3_ = 1000

*F*_*stall*_ = 0.86, *K*_*eff*0_ = 2, *K*_*eff*_ = 2, *δ*_6_ = 0.

Fig. 4G1

*K*_*s*_ = 6, *K*_*b*_ = 3, *K*_*BB*_ = 0, *F*_*cr*_ = 2, *d*_1_ = 1000, *d*_2_ = 1000, *d*_3_ = 1000

*F*_*stall*_ = 0.86, *K*_*eff*0_ = 2, *K*_*eff*_ = 2, *δ*_6_ = 0.

*K*_*s*_ = 3, *K*_*b*_ = 1.5, *K*_*BB*_ = 1000, *F*_*cr*_ = 1, *d*_1_ = .01, *d*_2_ = .01, *d*_3_ = .01

*F*_*stall*_ = 0.81, *K*_*eff*0_ = 1 to

### Numerical implementation for single cell behavior

Fig. 5C

*K*_*s*_ = 4, *K*_*b*_ = .4, *K*_*BB*_ = 0.25, *F*_*cr*_ = 0.4, *d*_1_ = .1, *d*_2_ = .1, *d*_3_ = .1

*F*_*stall*_ = 0.73

Fig. 5D

*K*_*s*_ = 4, *K*_*b*_ = .4, *K*_*BB*_ = 0.25, *F*_*cr*_ = 0.4, *d*_1_ = .1, *d*_2_ = 100, *d*_3_ = .1

Fig 5E:

*F*_*cr*_ = 0.4, *Soft gel*: *K*_*s*_ = 6, *K*_*b*_ = 1.0, *K*_*BB*_ = 0.6. *Stiff gel*: *K*_*s*_ = 60, *K*_*b*_ = 10, *K*_*BB*_ = 5, *F*_*cr*_ = 0.4

