## Supplementary material for "Opposing mechanical anchorage drives collective cell–matrix interactions": SI Figures

**Supplementary Information**

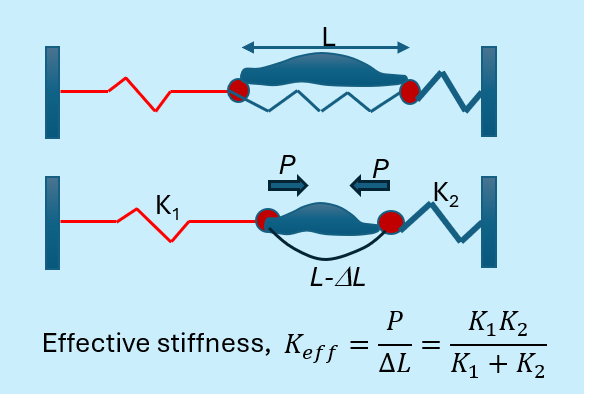

**SI Figure 1.** Mechanical analogue of a cell in ECM. The effective stiffness of the ECM felt by the cell against its force dipole is determined by the stiffness on both sides.

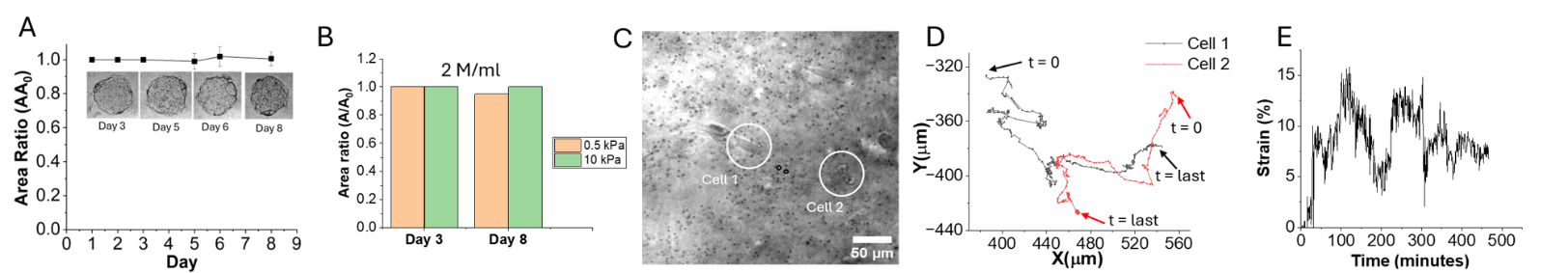

**SI Figure. 2.** Cells don’t compact linear elastic Col-T gels**.** (A) Snapshots of 0.5kPa Col-T gel discs with 3T3 cells (2 M/ml), and their normalized area ratio (A/A0) with time. (B) comparison in compaction between 0.5 kPa and 10 kPa a Col-T gel discs (2 M cells/ml), on day 3 and day 8. (C) Brightfield image of cells tracked with time (Fig. D). (D) Trajectory of 2 cells in (C) showing Brownian dynamics. (E) Strain measured in the region between cells 1 and 2 in C as a function of time using the displacements of 2 µm beads (dark spots) embedded in Col-T gel.

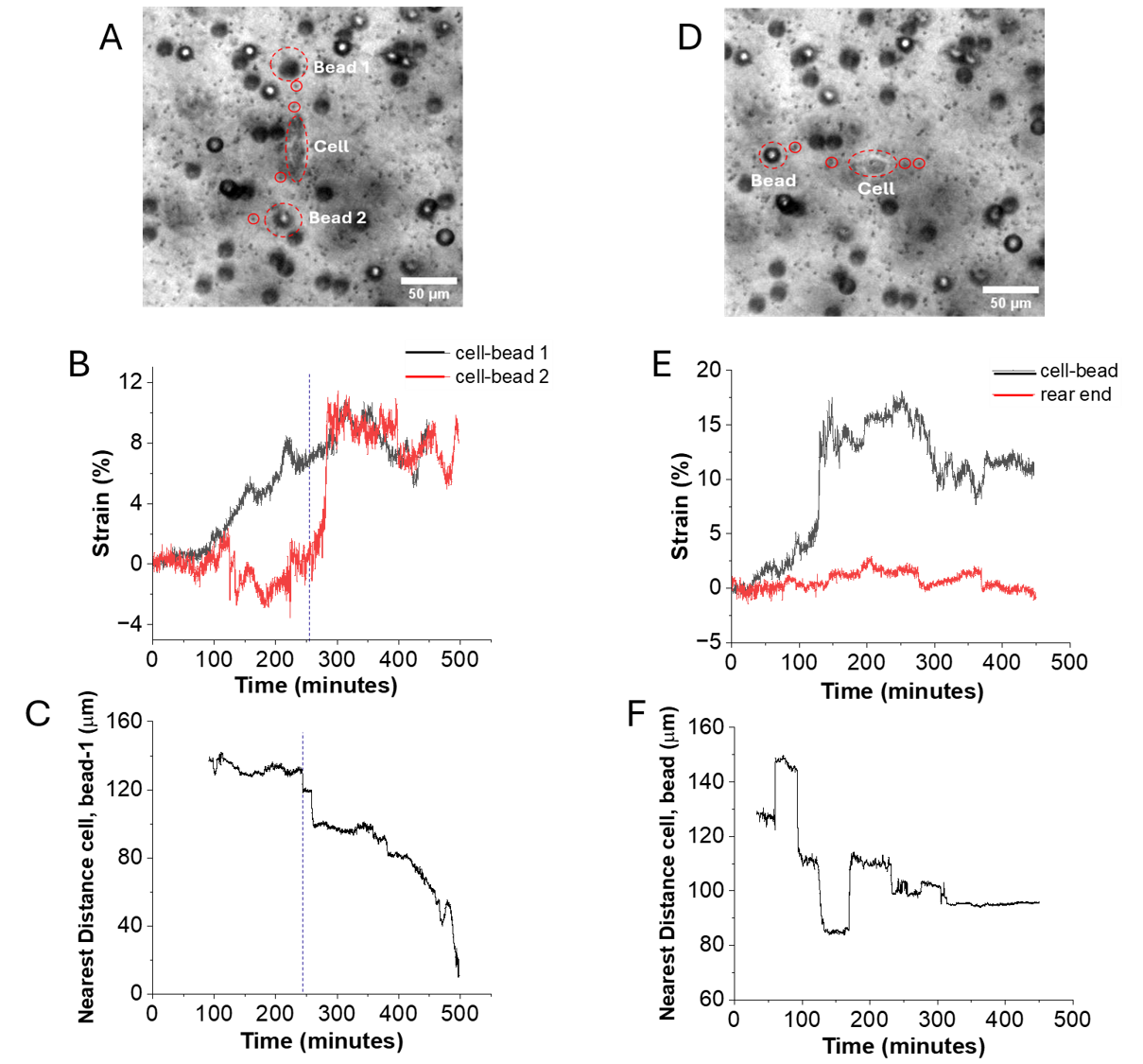

**SI Figure. 3.** Cells enter the feedback loop of strain escalation and persistent motion towards rigid beads (15 µm) when beads are present on opposite ends. (A) Brighfield image of a single cell with nearby beads is imaged over time. It has beads along the vertical direction (marked by red dotted circles). (B) Strain as a function of time along the vertical direction at leading (black) and rear (red) end of the cell. Leading and rear end are defined as the direction of motion of the cell and the opposite direction, respectively. Both strains increase over time and reach ~10%.(C) The distance between the filopodia tip of the cell and the top bead (A) at the leading edge as a function of time. The distance remain constant until 250^th^ minute, then it gradually decreases to zero as the cell moves closer to the bead. This time corresponds to the time when the rear end strain in B also start to rise sharply. This is shown by the vertical dotted line in B,C. (D) Brighfield image of the same cell interacting with a neighboring bead in the horizontal direction (marked by red dotted circles). There is no bead present at the rear end within the same radius as the leading edge. (E) Consequently, the strain between the cell and the bead at the leading edge rises to 15%, whereas the strain at the rear end remains very low. (F) The distance between the filopodia tip of the cell and the bead shows no temporal correlation with the rise in strain as in B-C. The distance slightly varies over time as the cell moves towards and away from the bead.

Bounds of $K_{eff}$

Eqs 2-5 give the bounds of $K_{eff}$. Eqs 2 and 3 imply that $\frac{d\delta_{1}}{dx}\leq0$ for all $\delta_{2}$, noting that $F_{*}\geq0$. Eq. 5 implies that $K_{eff}$ maximizes when $\frac{d\delta_{1}}{dx}=0$, i.e., $K_{eff\_max}=K_{s}$ as the upper bound. When $F_{*}=0$, $K_{eff}=\frac{K_{s}K_{b}}{K_{sb}}$. $K_{eff}$ is thus bounded by ($\frac{K_{s}K_{b}}{K_{sb}}$, $K_{s}$), accounting for all possible interactions with neighbors.

**Single cell behavior: predictions from theory**

Single cell behavior can be predicted using the 3-cell model using large cell-cell distance ($d_{i}=1000)$, and $K_{BB}=0.$

We consider two cases:

(1) Cell is far from any rigid boundary.

Soft matrix: cell remains in stochastic mode. We choose $\frac{K_{eff0}}{K_{0}}=1$. Corresponding $F_{stall}=0.63$. For $F_{cr}=1$, theory predicts that the cell force (normalized) $F_{c}(t)/F_{cr}$ remains stochastic and there is no stiffening of the matrix, i.e., $\frac{K_{eff}\left( t \right)}{K_{eff0}}=1,\forall t$ (Fig. 4E1).

Model parameters (Fig. 4E1):

$K_{s}=3, K_{b}=1.5, K_{BB}=0, F_{cr}=1, d_{1}=1000, d_{2}=1000, d_{3}=1000$

Calculated values: $F_{stall}=0.63, K_{eff0}=1, K_{eff}=1, \delta_{6}=0.$

### Stiff matrix: cell transitions from stochastic to persistent force mode*.* Here, $\frac{K_{eff0}}{K_{0}}=2$. Corresponding $F_{stall}=0.86$. For $F_{cr}=1$, theory predicts that cell force, $F_{c}(t)$, transitions from stochastic to persistent $F_{stall}$ (Fig. 4F1). The cell is expected to polarize and remain stationary. High matrix stiffness thus promotes cell polarization where cells can reach stall force, irrespective of whether the gel is fibrous or linear elastic.

Model parameters (Fig. 4F1):

$$K_{s}=6, K_{b}=3, K_{BB}=0, F_{cr}=1, d_{1}=1000, d_{2}=1000, d_{3}=1000$$

Calculated values: $F_{stall}=0.86, K_{eff0}=2, K_{eff}=2, \delta_{6}=0.$

### For cells with higher intrinsic critical force ($F_{cr}=2$), the cell remains in the stochastic mode, since $F_{stall}$ is much smaller ($F_{stall}$=.86) (Fig. 4G1).

Model parameters (Fig. 4G1).

$$K_{s}=6, K_{b}=3, K_{BB}=0, F_{cr}=2, d_{1}=1000, d_{2}=1000, d_{3}=1000$$

Calculated values: $F_{stall}=0.86, K_{eff0}=2, K_{eff}=2, \delta_{6}=0.$

(2) Cell with nearby rigid boundaries may self polarize. Cells between opposing rigid boundaries can be simulated using the 3-cell model with very stiff $K_{BB}$ and small cell-cell distance, $d_{i}$. Cells in soft matrix with $\frac{K_{eff0}}{K_{0}}=1$, $F_{stall}=0.63$ and $F_{cr}=1$, but $K_{BB}=1000$, and $d_{i}=0.1$. Theory predicts that cell force, $F_{c}(t)$, transitions from stochastic to persistent $F_{stall}$ when $K_{eff}$ increases from $K_{eff0}=1$ to a steady value (Fig. 4H1).

$K_{s}=3, K_{b}=1.5, K_{BB}=1000, F_{cr}=1, d_{1}=.01, d_{2}=.01, d_{3}=.01$

$Calculated values: F_{stall}=0.81, K_{eff0}=1$ to

**Multiple cell behavior: predictions from theory**

Model predicts gel compaction as a function of cell density: We vary cell-cell distance ($d_{1}=d_{2}=d_{3}$) from .1 to 5, with $F_{cr}=0.4$, and $K_{s}=4$, $K_{b}=0.4.$ Cell force, $F_{c}$, effective stiffness due to strain stiffening, $K_{eff}/K_{eff0}$, compaction, $\delta_{6}$, and cell-cell interaction force, $F_{*}$, are shown as a function of time in Fig. 5C. At high cell density or small cell-cell distance ($d_{i}$ = 0.1), cell force is stochastic for a short time. Cell-cell interaction then emerges as evidenced by increase in $K_{eff}$ (strain stiffening) and rise of cell-cell force, $F_{*}$. The cells soon reach a steady state with stall force. Steady compaction of the gel is similar for $d_{i}=0.1$ to 1 ($\delta_{6}\to1)$ (Fig. 10E) as observed in experiments (25) where all gel samples compact by 80% for all cell densities of 0.1 to 5 M/ml. This is a consequence of strong stiffening of the ECM under large deformation after buckling, and is mimicked in our model by the compressive spring $K_{BB}$. For cell densities between $d_{i}=1$ to 2, compression decreases sharply (Fig. 10C), also mimicking experimental observation. Beyond, $d_{i}=3$, there is no compaction.

Gel compaction needs at least 3 nearby cells: We apply the model with $d_{1}=0.1, d_{2}=100, d_{3}=0.1$, i.e., with two nearby cells, but the 3^rd^ is far away. The model predicts no cell-cell interactions and no compaction (Fig. 5D). Cell force is stochastic. Keff remains at the initial value of Keff0, and $F_{*}\left( t \right)=0$ for all time implying no cell-cell interaction. This prediction is consistent with experimental observation of two cells coming in close proximity in collagen gel, but migrating away without inducing any strain stiffening (Fig. 3B).

High cell density may not necessarily lead to gel compaction. Cell-cell interaction in 3D gel depends of cell type, gel initial stiffness, cell density, ECM mechanical properties. It is not feasible to test experimentally the combinations of a wide range of conditions. The model can simulate various conditions and reveal novel insights. Recall that the model predicts transition in single cell behavior from stochastic to steady state in stiff gels. We thus asked whether cells at high density in stiff matrix will interact with its neighbors and compact the gel. To test the possibility, we used the model to predict compaction and strain stiffening in a stiff gel (Ks=60, Kb=10, and KBB =5) and soft (Ks=6, Kb=1, and KBB =.5) matrix with ($d_{i}=0.05-5$, high to low density, Fig, 5E). $F_{cr}=0.4$ was applied for both cases. Soft gels compact significantly more than stiff gel at all cell densities. For example, at $d_{i}=0.1$, compaction is almost 100% and 5% in soft and stiff cases respectively. More importantly, the increase in effective stiffness due to strain stiffening and cell-cell interaction, measured by the ratio$K_{eff}/K_{eff0}$ after the cells have reached a steady state reveal that cells do not interact in stiff gel.

Fig. 5C

$$K_{s}=4, K_{b}=.4, K_{BB}=0.25, F_{cr}=0.4, d_{1}=.1, d_{2}=.1, d_{3}=.1$$

$$F_{stall}=0.73$$

Fig. 5D

$$K_{s}=4, K_{b}=.4, K_{BB}=0.25, F_{cr}=0.4, d_{1}=.1, d_{2}=100, d_{3}=.1$$

Fig 5E:

$$F_{cr}=0.4, {Soft gel: K}_{s}=6, K_{b}=1.0, K_{BB}=0.6. Stiff gel:K_{s}=60, K_{b}=10, K_{BB}=5, F_{cr}=0.4$$
